# Albumin-binding recombinant human IL-18BP ameliorates macrophage activation syndrome and atopic dermatitis via direct IL-18 inactivation

**DOI:** 10.1101/2023.05.30.542831

**Authors:** Young-Saeng Jang, Kyungsun Lee, Mihyun Park, Jin Joo Park, Ga Min Choi, Chohee Kim, Shima Barati Dehkohneh, Susan Chi, Jaekyu Han, Moo Young Song, Yong-Hyun Han, Sang-Hoon Cha, Seung Goo Kang

## Abstract

Given the clinical success of cytokine blockade in managing diverse inflammatory human conditions, this approach could be exploited for numerous refractory or uncontrolled inflammatory conditions by identifying novel targets for functional blockade. IL-18, a pro-inflammatory cytokine, is relatively underestimated as a therapeutic target, despite accumulated evidence indicating the unique roles of IL-18 in acute and chronic inflammatory conditions, such as macrophage activation syndrome. Herein, we designed a new form of IL-18 blockade, i.e., APB-R3, a long-acting recombinant human IL-18BP linked to human albumin-binding Fab fragment, SL335, for extending half-life. We then explored the pharmacokinetics and pharmacodynamics of APB-R3. In addition to an extended serum half-life, APB-R3 alleviates liver inflammation and splenomegaly in a model of the macrophage activation syndrome induced in IL-18BP knockout mice. Moreover, APB-R3 substantially controlled skin inflammation in a model of atopic dermatitis. Thus, we report APB-R3 as a new potent IL-18 blocking agent that could be applied to treat IL-18-mediated inflammatory diseases.

## Introduction

IL-18 is a potent pro-inflammatory cytokine that belongs to the IL-1 family of cytokines (1, 2). It was first well described for its role in inducing IFN-γ production from T cells and enhancing natural killer (NK) cell cytotoxic activity, thus conferring host defense against intracellular microbes (3-5). In addition to these cells, other immune and non-immune cells have been shown to express the IL-18 receptor (6-8), indicating the pleiotropic activity of IL-18 and, particularly, its potential contribution in inflammatory and autoimmune illnesses. Diverse immune-related diseases, such as type I diabetes (9, 10), multiple sclerosis (11, 12), rheumatoid arthritis (13), inflammatory bowel disease (14, 15), and adult-onset Still’s disease (AOSD) (16, 17), have been associated with IL-18 and their underlying pathophysiology has received considerable attention. Moreover, IL-18 knockout mice exhibited tolerance to disease induction, demonstrating less severe phenotypes than wild-type control mice in several inflammatory diseases, including experimental autoimmune encephalomyelitis, collagen-induced arthritis and type I diabetes model.

Patients with macrophage activation syndrome (MAS) exhibit high blood levels of IL-18, implying that IL-18 also plays a role in the pathogenesis of MAS (18). Recently, IL-18 was shown to be a key mediator of MAS in both IL-18BP knockout mice and NLRC4 gain of function mutant mice in that high levels of active IL-18 stimulates the production of other pro-inflammatory cytokines and promoting the excess activation of macrophages (19, 20). Importantly, blocking IL-18 has shown promise as a treatment strategy for MAS (19). In addition, IL-18 levels were found to be elevated in the skin and blood of patients with atopic dermatitis (AD) (21, 22). Notably, IL-18 promotes the Th2 immune response, including the induction of IL-4 and IL-13, and IgE production, which contribute to the development of AD (5, 23, 24). Thus, the balance between IL-18 and IL-18BP seems to be critical for maintaining immune homeostasis and preventing the development of inflammatory diseases, implicating that IL-18 is indeed a potential therapeutic target for diverse inflammatory disorders

Consequently, studies have explored the possibility of targeting IL-18 for disease management. Reportedly, anti-IL18 antibodies or IL-18 BP administration could ameliorate severity in several disease models. GSK 1070806, a humanized monoclonal antibody (mAb) against soluble IL-18, is currently under development for Crohn’s disease, AD, and Behcet’s disease (NCT03681067, NCT04975438, NCT03522662). Unlike anti-IL-18 mAb, IL-18BP is a natural antagonist of IL-18 capable of maintaining homeostasis of IL-18-mediated immune regulation. In murine models of inflammation, such as dextran sulfate sodium (DSS)-induced colitis, recombinant IL-18 BP could reduce disease severity (25, 26). In a model of rheumatoid arthritis, modest doses of IL-18 BP could successfully suppress inflammation (27). Consequently, Tadekinig alfa, a recombinant human IL-18BP, is under development for cytokine release syndrome, hemophagocytic lymphohistiocytosis (NCT05306080) and AOSD (28).

Based on the accumulated evidence, we aimed to design a more potent IL-18 drug candidate for IL-18-mediated inflammatory disease by extending the half-life of recombinant hIL-18 through SL335 fusion. SL335 is a Fab fragment of human anti-serum albumin antibody that possesses high binding affinity (K_D_: ∼1nM) to human serum albumin, and thus SL335 has a longer duration than control Fab in serum albumin through FcRn mediated recycling mechanism (29, 30). SL335 has been shown to exhibit a half-life of approximately 37 hours in rats, 10 times longer than the control common Fab fragment (29). Importantly, many functional recombinant proteins or ScFv can be fused to SL335, retaining their own functions and affording extended half-life *in vivo* (31). In the present study, we aimed to validate whether the SL335-fused human recombinant IL-18BP could induce potent therapeutic IL-18 blockade by undertaking pharmacokinetic (PK) studies in healthy mice, rats, and cynomolgus monkeys and pharmacodynamics (PD) studies in both MAS and AD model mice.

## Materials and Methods

### APR-R3 cloning and establishment of stable cell line

The codons of APB-R3 heavy and light chain genes, including signal peptide sequence, were optimized and cloned into a pCGS3 vector (ATUM, Vector ID: 226214). A stable cell line expressing APB-R3 was established in CHOZN^®^ ZFN-modified GS^-/-^ cell line (Sigma-Aldrich Co. LLC, US). Five research cell bank candidates were selected and stored through single-cell cloning. Finally, 1 master cell bank candidate among five research cell bank was selected and prepared at the GMP facility. The APB-R3 from the master cell bank was used for all the following studies.

### APB-R3 production

APB-R3 was obtained from a 14-day fed-batch culture and a three-step purification process. Briefly, the 1 vial of APB-R3 master cell bank was thawed and sequentially expanded from flask level and 50 L stirred bioreactor to obtain adequate cell numbers for 500 L main culture. The 14-day fed-batch culture was performed in a 500 L stirred tank bioreactor. On day 14, the cell culture fluid was filtered through two serialized depth filters (Merck Millipore, US), followed by 0.2 µm sterilized filtration (Sartorius, Germany). The purification process consisted of three-step chromatography, i.e., affinity chromatography using CaptureSelect^TM^ CH1-XL resin (Thermo Fisher Scientific, US) for capturing APB-R3, cation exchange chromatography using Fractogel^®^ EMD SO3-(S) resin (Merck Millipore, US) for intermediated purification, and anion exchange chromatography using Eshmuno^®^ Q resin (Merck Millipore, MA, US) for polishing. After the concentration and formulation step, APB-R3 purity was evaluated, and APB-R3 with ≥98% purity was used for all experiments.

### Animals

Using a C57BL/6 background, MACROGEN (Republic of Korea) generated IL-18BP knockout mice using CRISPR/Cas9 technology; IL-18BP KO mice were maintained in Kangwon National Universtiy under the approval of Institutional Animal Care and Use Committee (IACUC) (Study No: KW-210317-7) and transferred to QuBEST BIO Co., Ltd. (Republic of Korea) and used for CpG-induced mouse model study uner the approval of QuBEST BIO’s Institutional Animal Care and Use Committee (IACUC) (Study No.: 0903122160). SKH-1 hairless mice were purchased from Japan SLC, Inc (Japan). The mice experiments were performed under the approval of IACUC of Kyungpook National University for AD (Study No.: IV-210726-1.1). For PK studies, C57BL/6 mice and Sprague Dawley rats were maintained by KNOTUS (Republic of Korea), while cynomolgus monkeys were maintained by AnaPath GmbH (Switzerland), under standard care and approval.

### Evaluation of APR-R3 binding affinities

APB-R3 interactions with human serum albumin or IL-18 were determined via surface plasmon resonance using Biacore^TM^ T200 (Cytiva, US). After activation of the chip surface, APB-R3 (40 μg/mL) was flown over the chip surface at a flow rate of 30 μL/min for 120 s for immobilization, followed by washing and deactivation of excess APB-R3. To analyze the binding between APB-R3 and IL-18 or human serum albumin (HSA), IL-18 or HSA in a running buffer was flown over the chip at a rate of 30 μL/min for 120 s. For molecular dissociation, the washing buffer was flown over the chip at 30 μL/min for 2,000 s. Data were analyzed using Biacore^TM^ T200 software (Cytiva, US).

### In vitro bioassay of APB-R3 for IL-18 inhibition

The inhibitory effect of APB-R3 on IL-18 signal transduction was tested using the HEK-Blue^TM^ IL-18 reporter cell assay system (InvivoGen, US). APB-R3 and recombinant IL-18BP were appropriately diluted, mixed with IL-18 (1 ng/mL), and incubated at 37℃ for 30 min. For the assay, mixed samples were added to HEK-Blue^TM^ IL-18 reporter cells (5 × 10^4^ cells/well), followed by incubation at 37℃ for 21 h. Then, 40 μL of supernatants were used for the reporter assay, according to the manufacturer’s protocol. Human macrophage KG-1 cell line (ATCC, US) was used to test the inhibitory effect of APB-R3 on IFN-γ secretion. APB-R3 and recombinant IL-18BP were appropriately diluted, mixed with IL-18 (8 ng/mL), and incubated at 37℃ for 1 h. After the reaction, mixed samples were applied to KG-1 cells (1.3 × 10^5^ cells/well), followed by incubation at 37℃ for 23 h. The IFN-γ level in supernatants was quantitated using an ELISA kit (BioLegend, US) according to the manufacturer’s protocol.

### PK evaluation of APB-R3

The PK study using healthy Sprague Dawley rats was performed by KNOTUS Co., Ltd. (Republic of Korea) to compare the half-life of APB-R3 and recombinant IL-18BP. Briefly, healthy Sprague Dawley rats (males; n=3/group) were administered 2 mg/kg APB-R3 intravenously and 6 mg/kg APB-R3 subcutaneously. Meanwhile, recombinant IL-18BP was administered at 1 mg/kg via intravenous injection and 3 mg/kg via subcutaneous injection. Blood was collected and prepared for ELISA analysis at designated time points. For establishing the PK parameters in non-human primates (NHP) according to the administration route, a PK study was performed at AnaPath GmbH (Basel, Switzerland). Briefly, healthy cynomolgus monkeys (males, n=3/group) were administered 3 and 10 mg/kg APB-R3 via intravenous injection and 10 and 20 mg/kg APB-R3 via subcutaneous injection, respectively. Blood was collected and prepared for ELISA analysis at designated time points. To evaluate the PK profile of APB-R3 within a predictable therapeutic window in the disease mouse model, a PK study was performed using C57BL/6 mice by KNOTUS Co., Ltd. (Republic of Korea). Only one intravenous injection was administered. APB-R3 was administered at 10 and 20 mg/kg (males, n=10), respectively. Blood was collected and prepared for ELISA analysis at designated time points. In all PK studies, the serum APB-R3 concentration was quantified using the ELISA method and the PK parameters were evaluated by Phoenix^®^ WinNonlin^®^ (Certara, US).

### CpG-induced MAS mouse model

Wild-type C57BL/6 and IL-18BP knockout mice were randomly divided into groups intraperitoneally injected (IP) every other day with CpG ODN (50 μg/head; BIONEER, Republic of Korea). APB-R3 and PBS were intraperitoneally injected daily, along with body weight measurements. For biochemistry analysis, blood was collected via the intraorbital vein on day 10. Subsequently, mice were euthanized, and the livers and spleens were harvested. The serum aspartate transaminase and alanine transaminase levels were analyzed using an AU480 Chemistry analyzer (Beckman Coulter, US). The ferritin and Interferon-γ level were analyzed using the respective ELISA kits (Ab157713, Abcam, UK for ferritin and MIF00, RD systems, US for IFN-γ) according to the manufacturer’s instructions. For histopathological analysis, paraffin sections (3–4 μm thick) were prepared and stained with H&E according to standard procedures.

### Induction of atopic dermatitis and clinical assessment of the degree of AD

Each mouse was sensitized with 100 μL of 3% oxazolone solution under anesthesia on day 0. From day 7 to day 27, mice were challenged with 100 μL of 0.2% oxazolone solution thrice weekly. Mice were assigned to eight groups and administered treatment from day 7 to day 27. Anti-IL4Rα Ab and anti-IL18 Ab were intraperitoneally administered twice weekly from day 7. APB-R3 was intraperitoneally administered every other day from day 7. Skin erythema, scaling, and skin thickness were observed, and a score from 0 to 4 was assigned according to the following criteria, and the sum was used as a clinical activity index. The cumulative skin score is the sum of the point of each criterion; Erythema score (0∼4), Scaling score (0∼4), and skin thickness score (0∼4).

### Histomorphological evaluation

After separating the dorsal skin tissue, paraffin sections were prepared and stained with H&E. Tissue sections were examined under a microscope after H&E staining. The tissue sections were scanned and stored with a digital scanner, and each section was captured at 200× magnification and evaluated. Epidermal thickness was measured by more than two independent researchers.

### RT-qPCR

Following animal sacrifice, dorsal skin was harvested, and transcripts in skin tissues were quantitatively analyzed by RT-qPCR. After RNA extraction, cDNA was synthesized. Then, mRNA expression levels of five target genes were analyzed using each probe: TSLP (Mm01157588_m1), IL-1β (Mm00434228_m1), TNF-α (Mm00443258_m1), IL-4 (Mm00445259_m1), IL-13 (Mm00434204_m1) and IFN-γ (Mm01168134_m1). Probes are purchased from Thermo Fisher Scientific (US).

### Statistical analysis

Data are illustrated using GraphPad Prism 8.0 (GraphPad Software, Inc., US). Statistical analyses were conducted using GraphPad Prism 8.0. Comparisons were performed using a two-tailed Student’s t-test, Mann-Whitney U-test (Non-parametric test), or a two-way ANOVA. A p-value of ≤ 0.05 was considered statistically significant.

## Results and Discussion

### APB-R3 comprises SL335-linked recombinant human IL-18BP with extended half-life

APB-R3 was designed to prolong the serum half-life of recombinant human IL-18BP *in vivo* by linking hIL-18BP to the C-terminus of the heavy chain of human serum albumin-binding Fab, SL335 (SAFA) via flexible linker (Fig 1A). After assessing the eligibility for production and purifying APB-R3 stably, surface plasmon resonance was perfomed to evaluate whether APB-R3 could retain binding activity to both hIL-18 and HSA. We found that the equilibrium-binding affinity (K_D_) of APB-R3 to hIL-18 and HSA were 60.9 pM and 16.8 nM, respectively (Fig 1B), thereby suggesting that APB-3 has a high binding affinity to both hIL-18 and HSA. To determine the half-life of APB-R3, we performed a PK study in rodents and NHP. The PK study employed healthy male Sprague Dawley rats and compared hIL-18BP and APB-R3 administered via two distinct routes, intravenous (IV) and subcutaneous (SC). It should be noted that APB-R3 was administered at a two-fold higher dose than hIL-18BP to adjust the equivalent molar ratio between APB-R3 and rhIL-18BP (APB-R3; 79 kDa, hIL-18; 45 kDa). The half-life (*t_1/2_*) of APB-R3 and rhIL-18BP were 21.8 and 6.5 h following IV administration, and 34.9 and 9.7 h following SC administration, respectively (Fig 1C). Accordingly, the *t_1/2_* of APB-R3 was approximately 3-fold longer than that of rhIL-18BP following both IV and SC administration in Sprague Dawley rats (Fig 1C). Overall, APB-R3 demonstrated long-acting behavior when compared with rhIL-18BP, regardless of the administration route.

**Figure 1.**
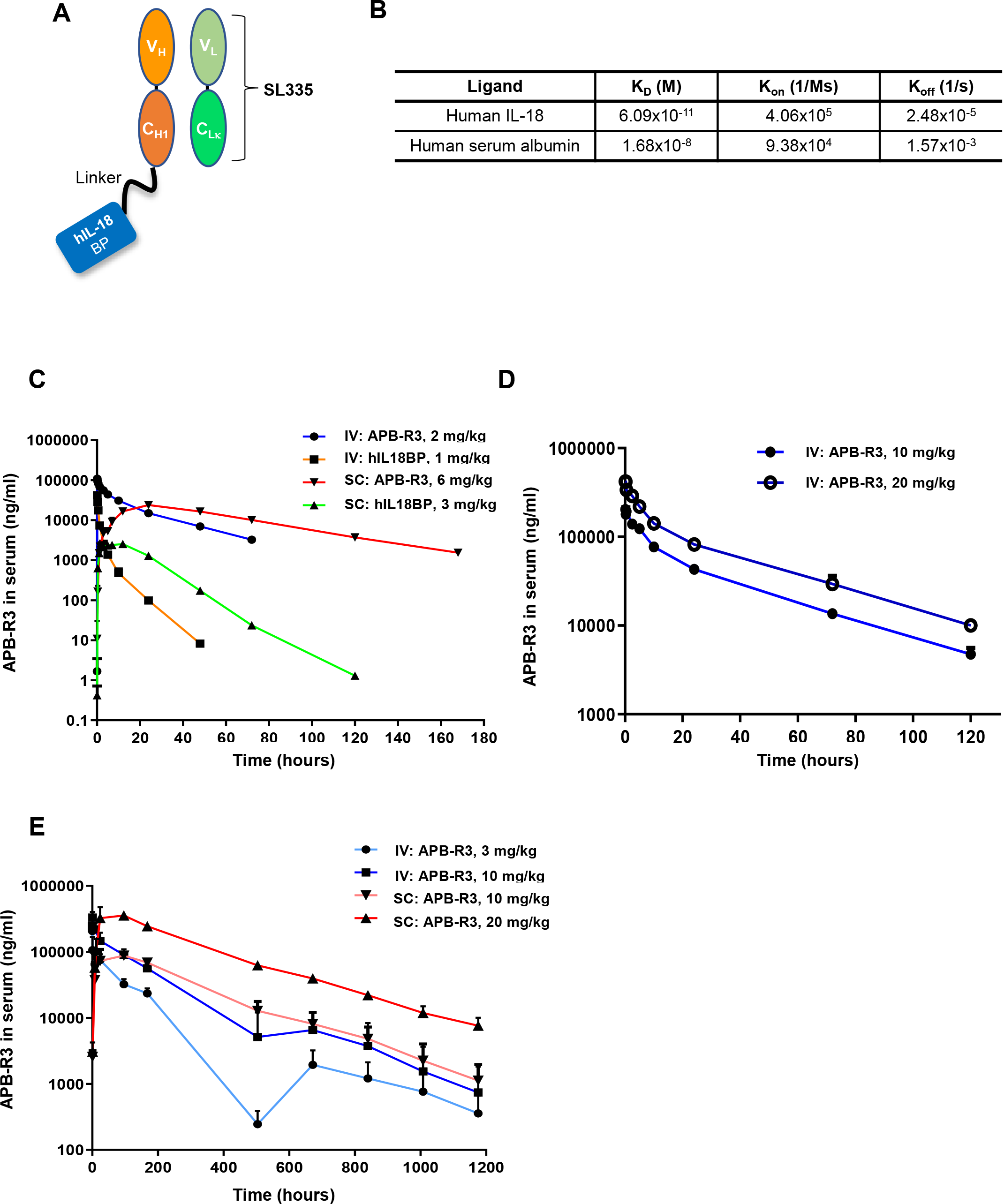
APB-R3 is a long-acting functional human recombinant IL-18BP. (A) The molecular structure of APB-R3. (B) Binding affinities of APB-R3 to hIL-18 and HSA were determined by surface plasmon resonance. (C-E). The PK profiles in rats, mice and cynomolgus monkeys. Serum APB-R3 levels were measured by quantitative PK ELISA at designated sampling time points, respectively (C) APB-R3 (2 and 6 mg/kg) and rhIL-18BP (1 and 3 mg/kg) were administered intravenously or subcutaneously to Sprague Dawley rats. (D) APB-R3 (10 and 20 mg/kg) was administered intravenously. (E) APB-R3 was administered intravenously (3 and 10 mg/kg) or subcutaneously (10 and 30 mg/kg) to monkeys. Data are presented as the mean ± SD.

Next, healthy C57BL/6 mice were used to evaluate the PK profile of APB-R3 to obtain a predictable therapeutic window in the mouse disease model for pharmacodynamic studies. Two different doses of APB-R3 (10 or 20 mg/kg, IV) were administered, and the PK profiles of APB-R3 showed dose-dependent behavior (Fig. 1D). Importantly, the *t_1/2_* of APB-R3 in mice was 30.2 h (10 mg/kg,) and 31.7 h (20 mg/kg). Thus, the *t_1/2_* of APB-R3 in mice was approximately 30 h, which was markedly similar to that in Sprague Dawley rats, revealing the long-acting behavior of APB-R3 in mice.

To evaluate the PK parameters of APB-R3 in NHP, an additional PK study was conducted in cynomolgus monkeys by administering APB-R3 via IV and SC injection, revealing *t_1/2_* values of 175.9 h (3 mg/kg, IV), 100.8 h (10 mg/kg, IV), 167.8 h (10 mg/kg, SC), and 206.6 h (20 mg/kg, SC) respectively (Fig. 1E). Although the *t_1/2_* of one group (10 mg/kg, IV) was relatively shorter than that of other groups, we concluded that the *t_1/2_* of APB-R3 in cynomolgus monkeys was approximately 7 days. Notably, human serum albumin-binding Fab, SL335, has been developed to target HSA and has a much higher binding affinity than rodent serum albumin (29). This property may contribute to the shorter half-life of APB-R3 in rodents than that in NHP. Nervertheless, our results clearly demonstrates the longevity of APB-R3 in both animal models which are comparable to those of native serum albumin considering that the the *t_1/2_* of serum albumin of rat is about 2 days and that of NHP. Overall, APB-R3 is a new IL-18 antagonist formulation with stable physicochemical properties and potentially long-lasting activity in humans.

### APB-R3 induces potent blockade of IL-18 signal transduction

APB-R3 was designed to achieve IL-18 blockade for therapeutic purposes. Hence, we aimed to determine whether APB-R3 could effectively inhibit IL-18 receptor signal transduction *in vitro*. Accordingly, the inhibitory effect of APB-R3 on the IL-18 signal transduction was measured using the HEK-Blue^TM^ IL-18 reporter cell assay system. APB-R3 efficiently blocked IL-18-induced signal transduction with similar potency to rhIL-18BP, indicating SL335 fusion did not induce steric hindrance (Fig 2A). Importantly, APB-R3 exhibited a superior inhibitory effect to that of the anti-IL-18 neutralizing antibody (EC_50_; 0.046 nM vs. 0.384 nM) (Fig 2A). It is well-established that macrophages also express functional IL-18 receptors, responsible for IFN-γ production under inflammatory conditions (6). Based on this cellular mechanism, the *in vitro* efficacy of APB-R3 was orthogonally evaluated using the human macrophage KG-1 cell line. Similar to the results of reporter cell assay, APB-R3 efficiently and dose-dependently inhibited IFN-γ secretion in the KG-1 cell line (Fig 2B), thereby supporting the possibility that APB-R3 could afford long-acting and functional blockade of IL-18 *in vivo*.

**Figure 2.**
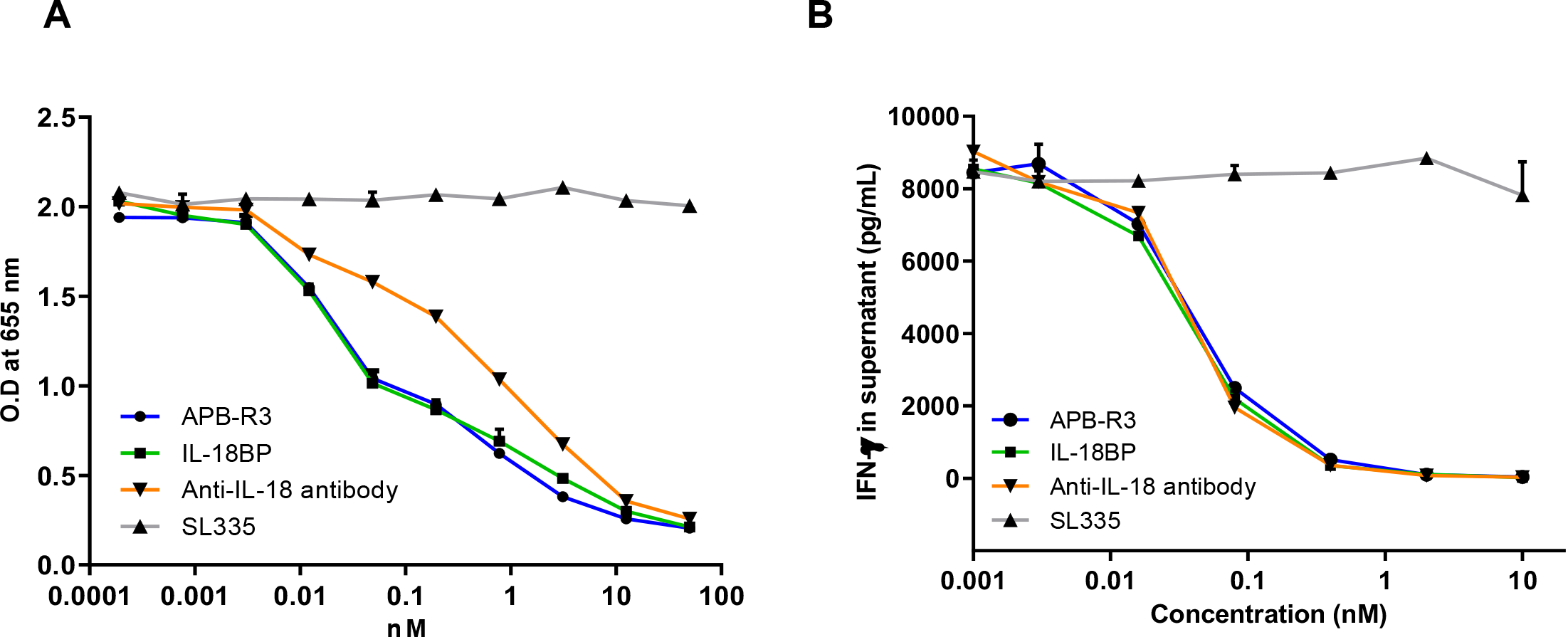
APB-R3 blocks IL-18R-mediated signaling activity. (A-B) *In vitro* bioassay to measure IL-18 induced signal transduction in HEKBlue^TM^ IL-18 reporter cell line (A) and IFN-γ production from KG-1 cell line (B). Cells were stimulated with hIL-18(1 ng/mL) and various amounts of IL-18 blockade, including the negative control, SL335. The experiment was performed in triplicate. Data are presented as the mean ± SD.

### APB-R3 ameliorates MAS symptoms induced in IL-18BP knockout mice by repeated CpG administration

To confirm the efficacy of APB-R3 in IL-18-related inflammatory model mice, IL-18BP KO mice were firstly generated by targeting *Il-18bp* with CRISPR/Cas9 technology in a C57BL/6 background embryonic stem cells, and the targeting was confirmed by sequencing and qRT-PCR for *Il-18bp* expression (Supplementary Fig. 1). Thereafter, a CpG-induced MAS mouse model was established using the IL-18BP knockout mice (19), and subsequently treated with two distinct doses of APB-R3 after MAS induction (Fig 3A). In the established model, acute severe inflammation primarily leads to liver dysfunction and induces splenomegaly (19); thus, we examined several parameters related to both organs. Our result showed that APB-R3 injections did not restore enlarged liver in MAS-induced mice. However, splenomegaly was dramatically restored in the APB-R3-treated group as demonstrated by relative weight to body weight (Fig. 3B). Treatment with low- (3 mg/kg) and high-dose (10 mg/kg) APB-R3 showed a tendency to restore levels of aspartate transaminase and alanine transaminase, although the levels were not statistically significant (Fig 3C). Importantly, APB-R3 resulted in low serum ferritin levels (Fig 3C), a well-known biomarker of MAS (32). The serum levels of IFN-γ, another biomarker of MAS (33), were markedly increased on day 4 and decreased on day 7 in the established inflammation model, and APB-R3 strongly suppressed IFN-γ secretion, maintaining stable levels during the study period (Fig 3D).

**Figure 3.**
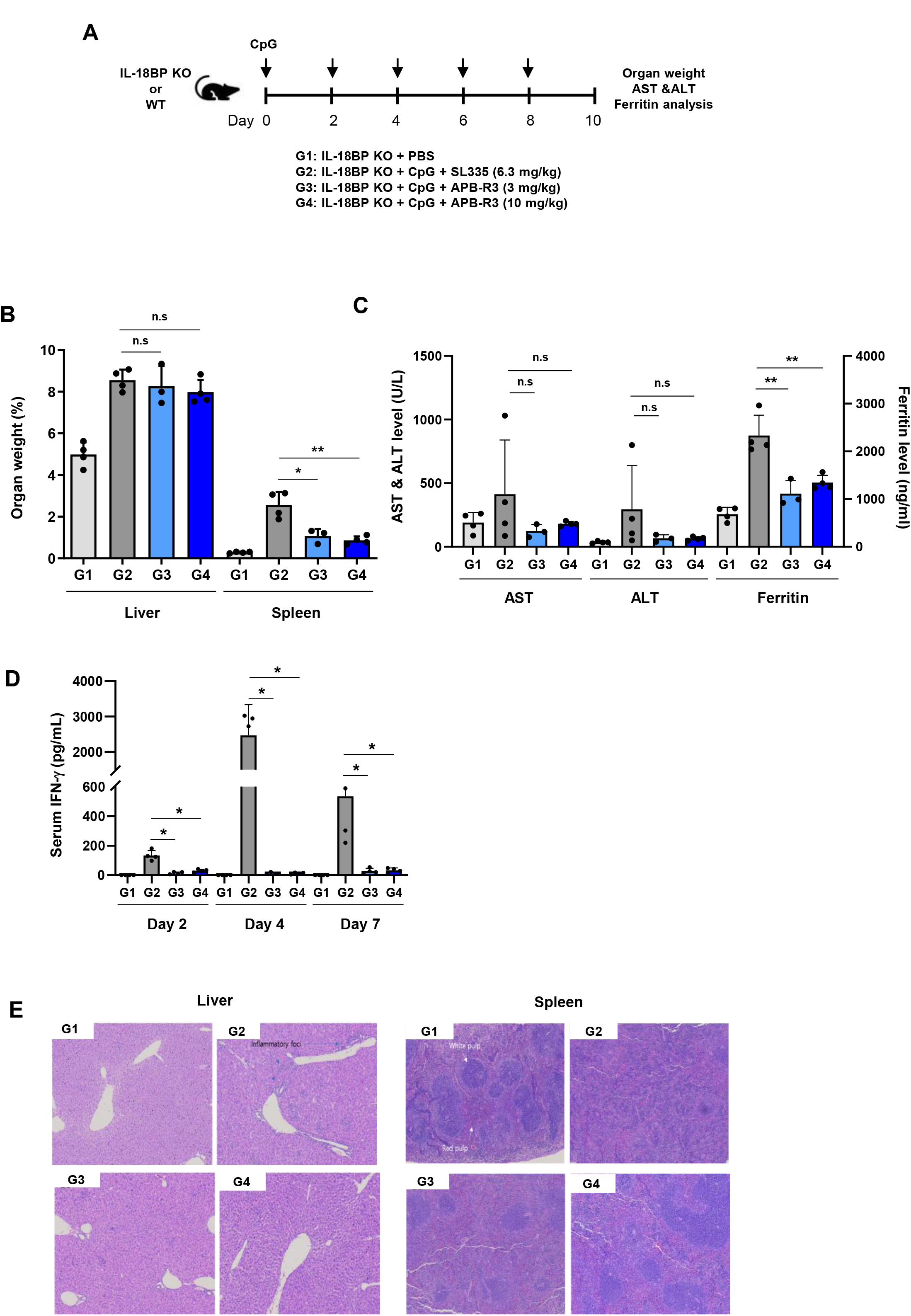
APB-R3 ameliorates liver toxicity and splenomegaly in MAS model mice. (A) A brief scheme to assess the efficacy of APB-R3 in the MAS mouse model. Five test groups were designated, as indicated. (B) The weight ratios of target organs per body weight were calculated by weighing animals and organs after sacrifice. (C) Aspartate transaminase, alanine transaminase, and ferritin levels were measured on the day of sacrifice. (D) Serum IFN-γ levels at each time point were determined by ELISA. (E) Representative images of H&E stained liver and spleen. Data are presented as the mean ± SD. Statistical analysis was performed using GraphPad Prism 8.0 (GraphPad Software, Inc., La Jolla, CA, USA). *p < 0.05, **p < 0.01.

In histopathological analysis, the APB-R3 treated group showed decreased gross inflammatory foci in the liver, accompanied by the restoration of disorganized white pulps in the spleen, a representative feature of severe acute inflammatory conditions, to a well-organized white pulp (Fig 3E). Moreover, multifocal mononuclear inflammatory cell infiltration and multifocal hepatocyte necrosis in the liver were reduced dose-dependently (Fig 3E). Taken together, these findings indicated that APB-R3 could efficiently prevent the inflammation-induced macroscopic lesion in the liver and spleen.

AOSD is a systemic autoinflammatory disease characterized by liver dysfunction, splenomegaly, lymphadenopathy, and eventually life-threatening conditions (34). Clinically, AOSD is similar to hemophagocytic lymphohistiocytosis, and MAS is both a common and substantial consequence of AOSD, with the level of IL-18 suggested as a precise diagnostic marker to distinguish between diseases (18). Reportedly, patients with AOSD exhibit considerably higher levels of IL-18 than patients with hemophagocytic lymphohistiocytosis and MAS (18). However, IL-18 is beyond a simple diagnostic marker. Instead, IL-18 could be a critical driver cytokine in AOSD pathogenesis, which is supported by the APR-R3-mediated resolution of clinical signs and reduction in serum IFN-γ and ferritin levels. However, other disease symptoms, such as weight loss, immune cell infiltration in the liver, and disorganized white pulp of the spleen, were poorly ameliorated by APB-R3 treatment, thereby indicating the need for a comprehensive understanding of specific lesions associated with immune regulatory pathways during uncontrolled inflammation in critically ill patients with MAS.

Overall, APB-R3 could afford potent IL-18 blockade, thereby preventing excess inflammatory response and ameliorating the IL-18-mediated pathological symptoms such as MAS, although we did not assess the long-acting potential of APB-R3 in this study.

### APB-R3 mitigates skin inflammation in oxazolone-induced AD

We evaluated the potential of APB-R3 in a CpG-induced MAS model in the presence of excess free active IL-18 owing to the IL-18BP knockout background. Most IL-18-related inflammatory diseases can be observed even in the presence of functional IL-18BP, and impaired IL-18BP-mediated inhibition of IL-18 activity is considered a key factor underlying disease pathogenesis. It is well-established that AD is an IL-18-related inflammatory disease that involves IL-4 and IL-13 expression (35). Thus, we aimed to establish the therapeutic potential of APB-R3 in oxazolone-induced AD, comparing the therapeutic potential of APB-R3 with that of IL-4 and IL-18 blockade using anti-IL-4R and anti-IL-18 neutralizing antibodies, respectively (Fig 4A). In oxazolone-induced AD, the skin appeared red and swollen on day 27, indicating well-established AD (Fig 4B). We observed that the gross skin lesion was markedly improved in all treatment groups, although only two APB-R3 doses, i.e., 3 and 10 mg/kg, afforded a statistically significant reduction in the cumulative skin score (Fig 4C). Low-dose APB-R3 (1 mg/kg) improved gross skin inflammation, as indicated by the area under the curve of the clinical score (Fig 4D). In addition, we quantitated erythema, scaling scores and skin thickness, and observed overall alleviation of skin inflammation in all treatment groups, with the APB-R3 treated groups showing a significant reduction when compared with other groups, especially the high-dose APB-R3 (10 mg/kg) group (Supplementary Fig. 2).

**Figure 4.**
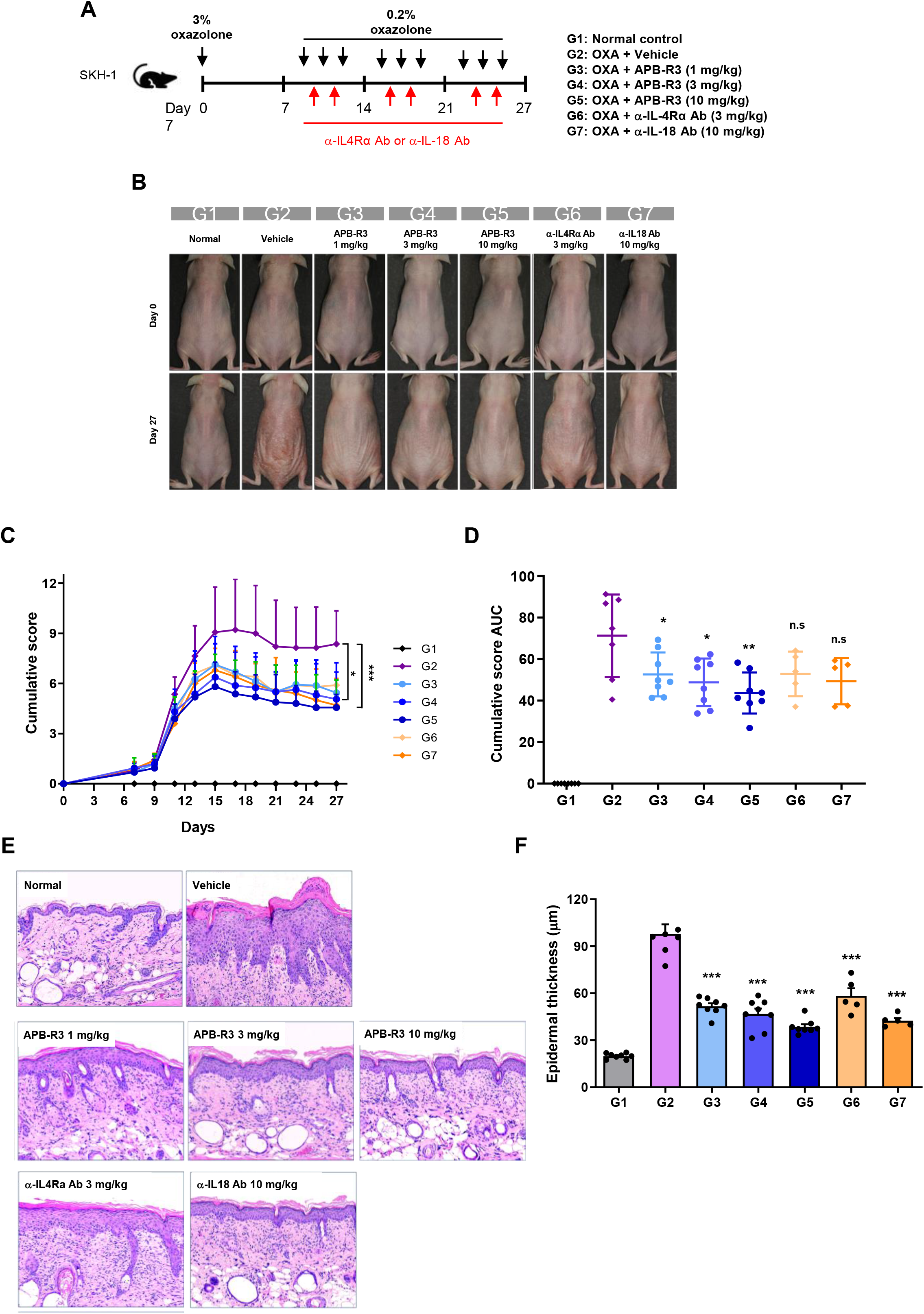
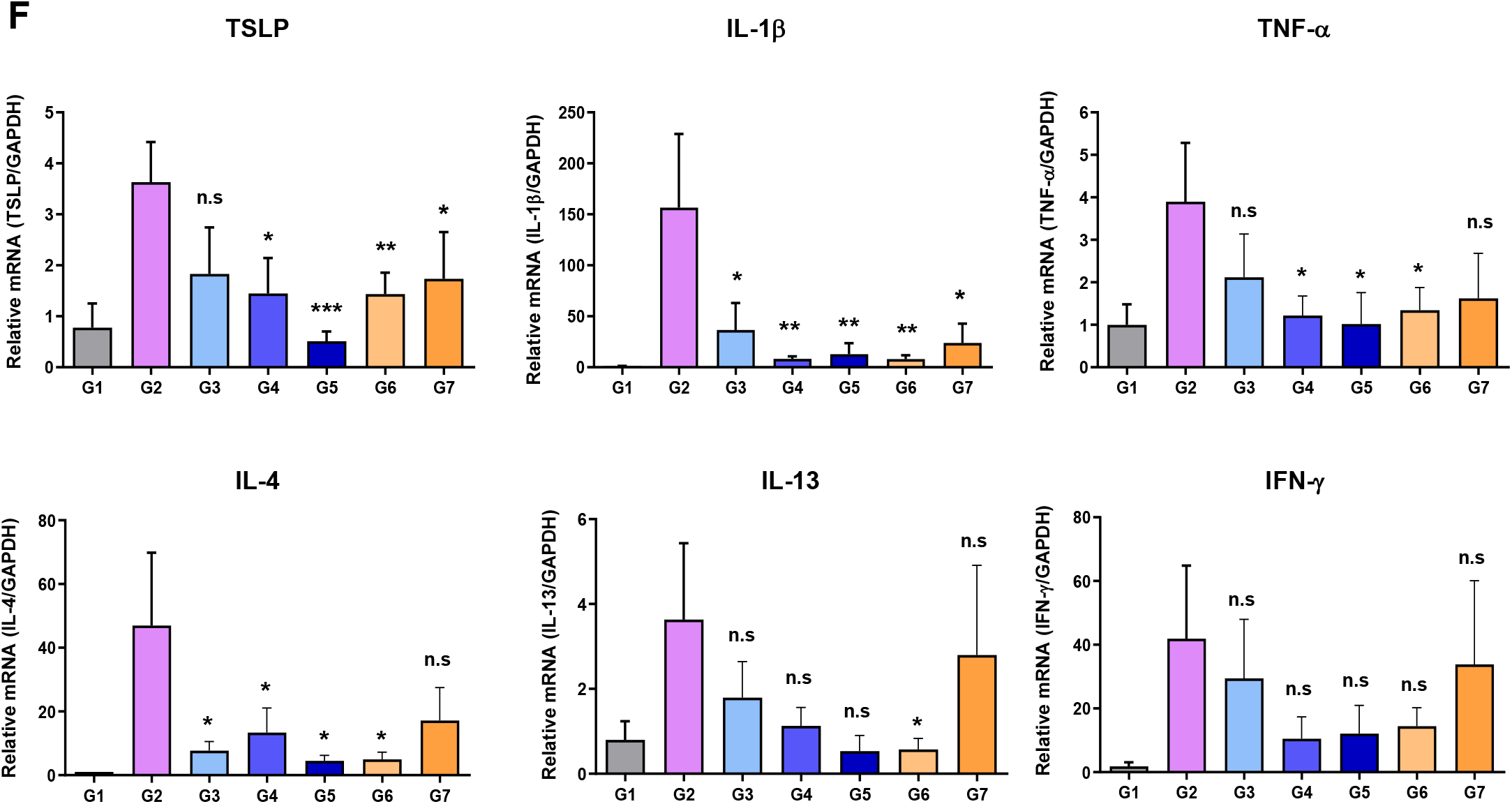
APB-R3 improves skin inflammation in the oxazolone-induced atopic dermatitis model. (A) A brief scheme to assess the efficacy of APB-R3 in oxazolone-induced atopic dermatitis (OIAD). Eight test groups were designated, as indicated. (B) Clinical features of OIAD mice following treatment with indicated compounds. (C) Cumulative score (erythema, scaling, and thickness) of each group of OIAD mice and AUC of cumulative score (D). (E) Histopathological features of skin lesions and (F) epidermal thickness of H&E stained skin tissues of each group of OIAD mice. (G) Relative mRNA expression level of TSLP, IFN-γ, IL-1β, TNF-α, IL-4 and IL-13 in skin tissues of oxazolone-induced atopic dermatitis mice, measured by RT-qPCR. Data are presented as the mean ± SD. *p < 0.05, **p< 0.01, and ***p < 0.001 versus vehicle control. Statistical significance was established using independent t-tests.

To comprehensively elucidate the effects of APB-R3 on skin inflammation, we performed histological analysis and measured the expression of inflammatory cytokines in skin tissues. Histological analysis revealed the thickened epidermal layer in control mice, which was alleviated in all treatment groups, including the low-dose APB-R3 (1 mg/kg) group (Fig 4E, F). In addition, all other histological features of AD, including dermal hyperplasia, spongiosis and hyperkeratosis in the epidermis, swelling, inflammatory cell infiltration, and enhanced angiogenesis and interstitial matrix, were observed and notably improved following treatment with APB-R3 (Fig 4E). Moreover, mRNA levels of TSLP, IL-1β, TNF-α, and IL-4 were significantly decreased in APB-R3 treated groups (3 and 10 mg/kg) (Fig 4G). Moreover, treatment with the lowest APB-R3 dose (1 mg/kg) could significantly reduce mRNA levels of IL-1β and IL-4 (Fig 4G). Treatment with APB-R3 also reduced mRNA levels of IFN-γ and IL-13, although no statistical significance was detected (Fig 4G). These results indicate that APB-R3 could suppress inflammation-mediated cytokine production in the skin, which might be responsible for the amelioration of gross skin inflammation, as determined by histology and clinical assessment. Based on the above results, APB-R3 showed a similar or even better therapeutic efficacy than anti-IL-4R or anti-IL-18 antibody in terms of clinical assessment. Moreover, treatment with high-dose APB-R3 (10 mg/kg) consistently demonstrated a better effect than anti-IL-18 antibody (Fig 4), thereby suggesting that APB-R3 is a potent therapeutic for blocking active IL-18 *in vivo*.

The IL-4 receptor alpha (IL-4Ra) antagonistic antibody, dupilumab, has been successfully developed for clinical application in AD, highlighting the importance of Th2-type inflammation in AD (36). It is well-established that IL-18 amplifies Th1-type immune responses by stimulating IFN-γ production. Moreover, a single nucleotide polymorphism (rs 187238) in the IL-18 locus was shown to be a genetic risk factor for AD (37). Elevated levels of IL-18 have been documented in the serum and skin of patients with AD (38, 39), indicating the presence of IL-18-mediated pathogenies in AD. Importantly, IL-18 also stimulates the production of Th2 cytokines, IL-4 and IL-13, by activating mast cells and basophils(5, 7). Therefore, it is plausible that IL-18 is an upstream trigger of both Th1- and Th2-type responses in AD pathogenesis, which could explain the superior efficacy of APR-R3 in the oxazolone-induced AD model, considering the cumulative score and histological assessment. Collectively, we demonstrated that APB-R3 could potently block IL-18 activity and thus abate IL-18-mediated inflammation in the AD mouse model, suggesting a potent candidate for further therapeutic development.

## Abbreviations

AD: atopic dermatitis
AOSD: adult-onset Still’s disease
HSA: human serum albumin
MAS: macrophage activation syndrome
OIAD: oxazolone-induced atopic dermatitis

## Acknowledgmentsfg

This work was supported by the National Research Foundation of Korea (NRF) grant funded by the Korea government (MSIT) (2021R1A2C1012421 to S.G.K, and 2021R1C1C1004023 and 2021R1A4A3031661 to Y.-H.H)

## Author Contributions

Y.-H.H., S.-H.C., and S.G.K. conceived and designed the project. Y.-S.J., M.P., C.K., performed and analyzed the experiments with under the supervision of Y.-H.H., S.G.K.. M.P., C.K., and S.B.D., bred and provided IL-18BP KO mice. J.H., G.M.C. produced APB-R3 including APB-R3 expressing cell line development with under the supervision of S.-H.C.. K.L., J.J.P., M.Y.S., S.C. supported the experiments performed by CRO with under the supervision of S.-H.C. and S.G.K.. Y.-S.J., K.L, and S.G.K. wrote the manuscript with input from the other authors. All authors contributed to the article and approved the submitted version.

## Figure legends

**Supplementary Figure 1.**
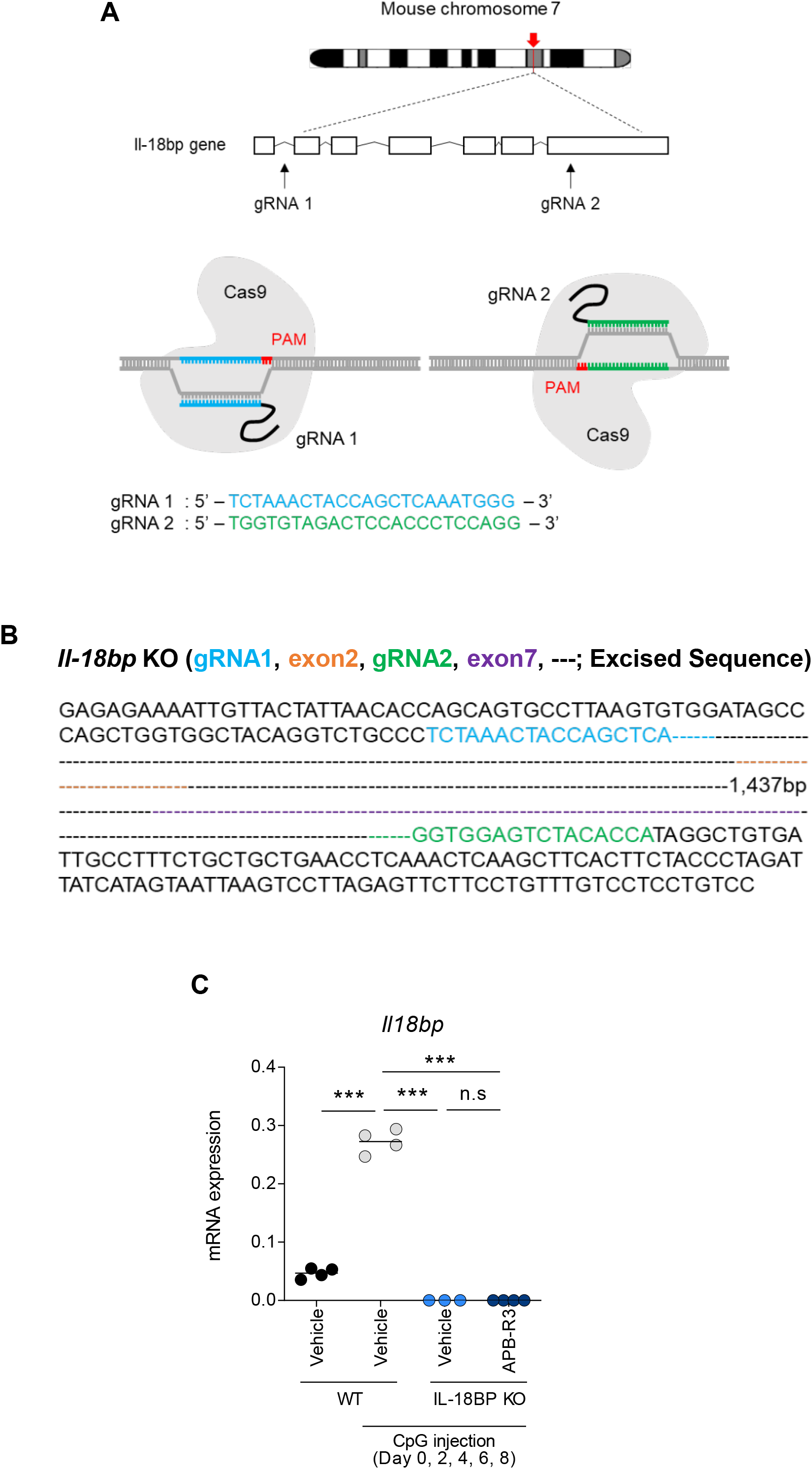
Generation of *Il-18bp* knockout mice and validation. (A) Strategy for targeting *Il-18bp* using CRISPR/Cas9. (B) Validation of *Il-18bp* modification via sequencing (C). Evaluation of IL-18BP mRNA expression in IL-18BP knockout mice in spleenocytes of MAS induced mice.

**Supplementary Figure 2.**
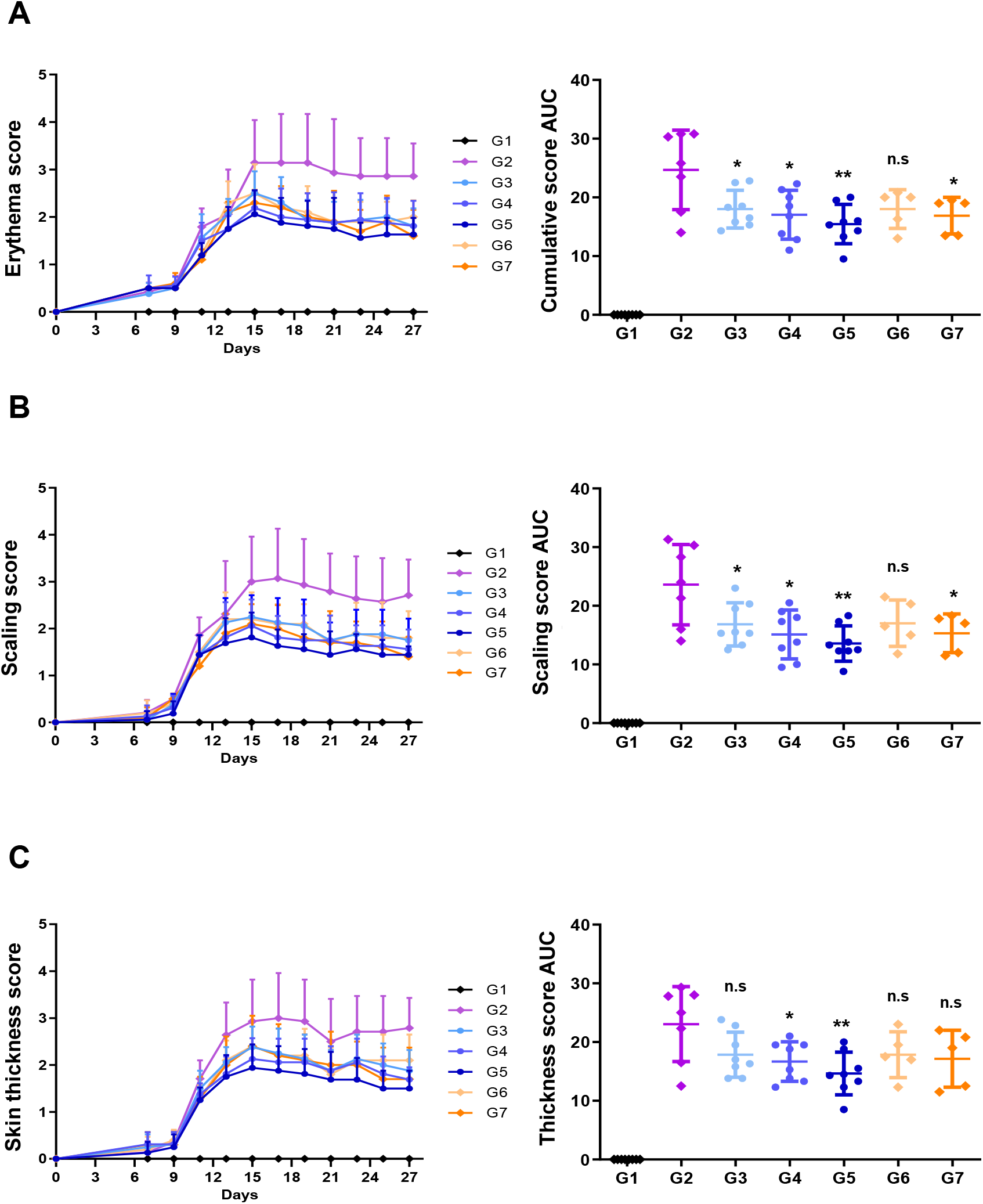
APB-R3 improves erythema and scaling in oxazolone-induced atopic dermatitis mice. (A-C) Experimental results of Figure 4. Erythema (A), scaling (B), and skin thickness (C) score of each group of oxazolone-induced atopic dermatitis mice. Data are presented as the mean ± SD. * p < 0.05, **p< 0.01, and ***p< 0.001 versus vehicle control. Statistical significance was established using repeated measures ANOVA.

